# Low microbial abundance and community diversity within the egg capsule of the oviparous cloudy catshark (*Scyliorhinus torazame*) during oviposition

**DOI:** 10.1101/2024.02.28.582473

**Authors:** Wataru Takagi, Ayami Masuda, Koya Shimoyama, Kotaro Tokunaga, Susumu Hyodo, Yuki Sato-Takabe

## Abstract

Vertebrate embryos are protected from bacterial infection by various maternally derived immune factors before the embryonic organs are fully developed. However, the defense mechanisms employed by elasmobranch embryos during development remain poorly understood. This study attempted to elucidate the embryonic defense mechanism of elasmobranchs by investigating the intracapsular environment of freshly laid eggs of the oviparous cloudy catshark (*Scyliorhinus torazame*). The egg capsule of oviparous elasmobranchs is tightly sealed until pre-hatching (early opening of the egg capsule), after which seawater flows into the capsule and the embryos are consequently exposed to the surrounding seawater. We first experimentally examined the resistance of embryos to potential bacterial infections and found that the early embryos were highly vulnerable to environmental pathogens, suggesting that the embryos are protected from the threat of infection before pre-hatching. Indeed, the intracapsular environment of freshly laid eggs exhibited a significantly low bacterial density that was maintained until pre-hatching. Furthermore, the microbiome inside eggs just after oviposition differed markedly from the microbiomes of rearing seawater and adult oviducal gland epithelia; these eggs were predominantly populated by an unidentified genus of Sphingomonadaceae. Overall, this study provides compelling evidence that early embryos of oviparous cloudy catshark are incubated in a clean intracapsular environment that potentially plays a significant role in embryonic development in oviparous elasmobranchs. Our results suggest that maintenance of this clean condition might be attributable to bactericidal or bacteriostatic activities associated with the egg jelly and/or oviducal gland.

## Introduction

Elasmobranchs (sharks, skates, and rays) exhibit remarkable diversity in reproductive strategies, ranging from oviparity to placental viviparity, accompanied by various modes of supplying embryonic nutrition (Wourms, 1977; Luer and Wyffels, 2022). Regardless of their reproductive mode, all elasmobranchs undergo internal fertilization via copulation, and their eggs are subsequently enclosed in a tertiary egg capsule, produced by the oviducal gland (OG, also known as the egg-shell or nidamental gland) (Hobson, 1930; Hamlett et al., 1998). In most oviparous species, the incubation period spans at least several months, during which embryos develop outside the mother’s body and rely solely on egg yolk for nutrition (Wourms, 1977; Luer and Gilbert, 1985; Ballard et al., 1993; Musa et al., 2018). In oviparous species, the inside of the capsule is filled with egg jelly surrounding the fertilized egg at oviposition. The jelly consists of three layers with distinct properties: a liquid layer closest to the embryo, a viscous colloid layer, and a semi-translucent solid layer (Wyffels et al., 2006). At oviposition, the solid jelly tightly seals both terminal ends of the capsule and then gradually liquefies as development progresses (Ballard et al., 1993). The anterior end of the capsule subsequently opens during the mid-period of development, an event known as “pre-hatching” or “eclosion”. Oviparous elasmobranchs produce tough and durable egg capsules, that protect the embryos against various challenges in the marine environment, such as mechanical stress and predation (Ballard et al., 1993; Kormanik, 1993; Musa et al., 2018). Before pre-hatching, the presence of a hole in the egg capsule wall of sufficient size to allow the influx of seawater will reportedly lead to the immediate death of the embryo (Ballard et al., 1993). In such cases, infection with marine pathogenic bacteria is considered the primary cause of embryonic death. Indeed, this hypothesis is supported by reports that the addition of antibiotics to rearing buffered-NaCl solution or seawater can extend the lifespan of early embryos outside the capsule in several oviparous shark species (Onimaru et al., 2018; Kuroda et al., 2021). However, when the embryos themselves acquire resistance to environmental pathogens and how they are immunologically protected before this acquisition remain largely unknown.

The capsule wall is highly porous, with an estimated pore radius of 13.6 Å (= 1.36 nm) in *Scyliorhinus canicula* (Hornsey, 1978), which is significantly smaller than the average size of a bacterium (Robertson et al., 1975). The capsule wall of oviparous cartilaginous fishes is highly permeable to small molecules such as dissolved gases, water, urea, electrolytes, and glucose (Hornsey, 1978; Kormanik, 1993; Takagi et al., 2014; 2017). In other words, the entry and exit of bacteria before the pre-hatching period are likely minimal, whereas small molecules are frequently exchanged across the capsule wall. Furthermore, the egg capsule itself reportedly exhibits anti-fouling properties (Thomason et al., 1994; 1996). These characteristics of the capsule wall are thus thought to prevent the invasion of pathogenic bacteria or fungi from the external environment.

Unlike the function of the capsule wall, details regarding the function of egg jelly in this context remain unclear and debatable. Physical protection of the embryo is considered the primary role of egg jelly (Koob and Straus, 1998; Musa et al., 2018). In other oviparous vertebrates, the substances surrounding the fertilized egg, such as hen’s egg white, contain an abundance of bactericidal proteins that contribute to defense against pathogens (Giansanti et al., 2012). Meanwhile, presence of antimicrobial substance has not been demonstrated thus far in the egg jelly of oviparous elasmobranchs (Lenain and Henderson, 2020). On the contrary, antibacterial activity against several gram-positive bacteria was observed in an investigation of the egg jelly from *Scyliorhinus stellaris*, suggesting the presence of antimicrobial substances (Martinengo et al., 2024).

Details regarding the bacterial flora, symbiotic interactions with the hosts, and types of pathogenic bacteria in elasmobranchs have only recently begun to be reported (Givens et al., 2015; Doane et al., 2017; 2020; Kearns et al., 2017; Pogoreutz et al., 2019; Mika et al., 2021; Storo et al., 2021; Muñoz-Baquero et al., 2023). A pioneering study of the oviparous little skate, *Leucoraja erinacea,* characterized the microbiota within the egg capsule and demonstrated intergenerational vertical microbial transmission (Mika et al., 2021). Although this previous work provided the first information regarding the microbiological environment surrounding oviparous embryos, how the bacterial flora within the capsule contributes to embryonic defense remains unclear.

Here, we experimentally examined the pathogen resistance of developing embryos and investigated the intracapsular microbial environment in the eggs of the cloudy catshark, *Scyliorhinus torazame*. The bacterial density within the egg capsule was directly measured, and a microbiome analysis was also conducted to characterize the microbial community.

## Methods

### Animals

Adult female catsharks were transported from the Ibaraki Prefectural Oarai Aquarium to the Atmosphere and Ocean Research Institute and kept in 1,000 and 3,000 L holding tanks with recirculating natural seawater at 16°C, maintaining a constant photoperiod (12 h light:12 h dark). The egg-laying cycle of catshark occurs approximately once every two or three weeks (Inoue et al., 2022). The fertilized eggs were manually collected from the cloaca of females, tagged to identify the date of oviposition, and subsequently placed in a separate tank away from adult individuals. Developmental stages were primarily identified according to the staging criteria of the closely related *S. canicula* (Ballarad et al., 1993). The occurrence of pre-hatching was visually determined by confirming whether air entered the capsules when taken out of water. Following the previous study (Honda et al., 2020), stage 31, the pre-hatching stage, was subdivided into stages 31E and 31L based on whether the eggs were pre-hatched. All animal experiments were approved by the Animal Ethics Committee of the Atmosphere and Ocean Research Institute of the University of Tokyo (P19-2). The present study was carried out in compliance with the ARRIVE guidelines.

### Survival test

Embryos just before pre-hatching (st. 30-31E) and earlier embryos (st. 27-28) were carefully removed from the egg capsules and individually transferred to 60 mL rearing containers, each filled with seawater under the following conditions. One was supplemented with an antibiotic-antifungal mixture (100x concentrated, penicillin 10,000 U/mL, streptomycin 10,000 µg/mL, amphotericin B 25 µg/mL, 0.85% saline, Nacalai Tesque, Kyoto, Japan) to achieve a final concentration of 0.2% (st. 31E, *N* = 6; st. 27-28, *N* = 6), while the other was without antibiotics (st. 31E, *N* = 5; st. 27-28, *N* = 5). The containers were maintained at 16°C in an air incubator, and survival days were recorded. The culturing seawater was changed weekly.

### Bacterial abundance

The surfaces of all the eggs and dissecting instruments were sterilized just before sampling using paper towels soaked in 70% ethanol. The anterior end of the egg capsules was trimmed with sterilized scissors, and the egg jelly within the capsule was aseptically collected using disposable pipettes. Embryos in normally developing eggs become observable one month after oviposition when exposed to intense light from the outside. Meanwhile, in unfertilized or developmentally arrested eggs, for some reason, the egg yolk swells, leading to irregular disintegration (Figure 1B). In this study, such eggs that did not progress in development even after a month were designated as “dead eggs”. Total contents inside the egg capsule containing egg jelly and yolk were collected from the dead eggs (*N* = 7), while only aqueous egg jelly surrounding the embryos was collected from the eggs containing embryo (st. 27-28, *N* = 6; st. 30-31E, *N* = 6). For the freshly laid eggs, only viscous jelly was collected immediately after oviposition and homogenized in a sterilized stomacher bag (Sansei medical Co. Ltd., Kyoto, Japan) using a rubber roller (*N* = 6). Each sample, whose density is assumed to be close to 1 kg/m^3^, was weighed and diluted with sterile distilled water to achieve a total volume of 1 mL. Undiluted 8 mL of seawater from the rearing container was also collected (*N* = 1). All samples were immediately fixed by adding 10% neutral buffered formalin (Fujifilm Wako Pure Chemical Corporation, Osaka, Japan) to achieve a final concentration of 1% in the dark at 4°C. Fixed samples were stained with 4′,6-diamidino-2-phenylindole (DAPI) for 10 minutes in the dark and then filtered through Nuclepore black polycarbonate membrane filters (0.22 µm pore size, Merck Millipore, Darmstadt, Germany) with negative pressure (≤ 20 kPa). The filtered membranes were mounted on glass slide, covered with a coverslip using immersion oil (IMMOIL-F30CC, Olympus, Tokyo, Japan), and gently pressed by thumbs to be flattened for observation. Under a fluorescent microscope, 20 random fields were selected from the entire membrane. In each field, the bacterial count within a defined area (2,000 µm^2^ for samples with high bacterial abundance and 10,000 µm^2^ for other samples) was measured, and the total bacterial count on the membrane was calculated. Eventually, considering the volume of the original sample and dilution factor, the bacterial density per unit volume (cells/mL) was determined. Differences in the density between groups were statistically analyzed by the Kruskal–Wallis test with GraphPad Prizm version 9.4.0 (San Diego, CA, USA).

**Figure 1:**
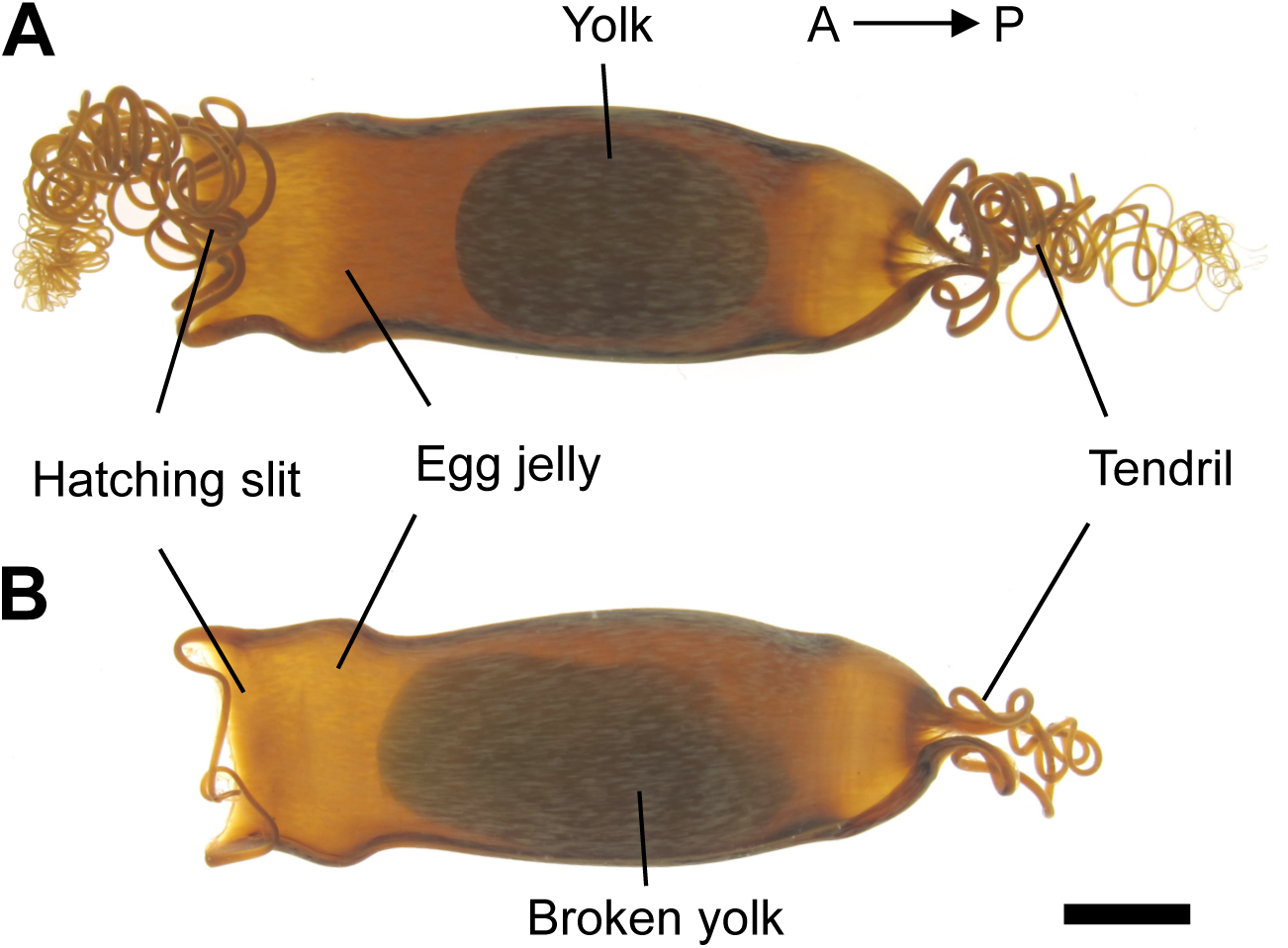
Eggs of cloudy catshark. The fertilized egg is encapsulated in the tough capsule filled with egg jelly. Arrows indicates the anterior-posterior axis with respect to the adult female. Prior to pre-hatching period (before stage 31E), the jelly plugs both the anterior and posterior ends of the capsule, preventing the entry of surrounding seawater. At pre-hatching, the anterior end of the capsule (hatching slit) opens. **A** Freshly laid egg. **B** “Dead” egg, containing yolk of which acellular vitelline membrane was broken. Scale bar, 1 cm.

### DNA extraction and 16S rRNA amplicon sequencing

Using the same procedure as mentioned above, egg jelly was collected from freshly laid eggs (*N* = 10) and dead eggs (1 month after oviposition, *N* = 6; 2.5 months after oviposition, *N* = 6). The sample of the dead eggs included egg yolk. Seawater (200 mL) from the container rearing eggs was also collected and filtered through a 0.22 µm filter (*N* = 1). The female individuals were anesthetized with 0.02% (w/v) ethyl 3-aminobenzoate methanesulfonate (Sigma-Aldrich, St. Louis, MO, USA), euthanized by decapitation, and dissected. The oviducal gland was carefully incised using sterile scissors. Nuclease-free water (Qiagen, Hilden, Germany) was then applied onto the inner epithelium of the gland, pipetted several times, after which the liquid was collected into microtubes. All samples were immediately snap-frozen in liquid nitrogen and stored at -30°C until use.

Total DNA was extracted using the DNeasy Blood & Tissue Kit (Qiagen) following the manufacturer’s instructions. The extracted DNA was subsequently sent to Genome– Lead Co., Ltd. (Kagawa, Japan) for library preparation, quality control, and sequencing, according to Illumina’s official protocol (Illumina, 2013). A first PCR reaction was performed with KAPA HiFi 2X Mastermix (KAPA) to amplify the V3-V4 region of 16S rRNA, using the following primer sets: forward primer: 5’-TCGTCGGCAGCGTCAGATGTGTATAAGAGACAGCCTACGGGNGGCWGCAG-3’, reverse primer: 5’-GTCTCGTGGGCTCGGAGATGTGTATAAGAGACAGGACTACHVGGGTATCTAA TCC-3’. The PCR procedure was as follows: initial denaturation at 95 °C for 3 min, 28-32 cycles of 95 °C for 30 sec, 55 °C for 30 sec, and 72 °C for 30 sec in a reaction volume 25 µl. The number of PCR cycles was determined based on the amplification efficiency. The PCR products were purified and barcoded with adapter and index sequences by a second PCR. The concentrations of the purified PCR products were normalized, and 301 bp × 2 paired-end reads were sequenced on the Illumina MiSeq platform. The acquired raw data were deposited in the NCBI Sequence Read Archive (Accession No: DRR523913-523937).

Community analysis of the reads was performed using the QIIME2 program (ver. 2023.9; Bolyen et al., 2019) as described below. First, primer sequences were removed using the Cutadapt plug-in (Martin, 2011), and the sequence reads potentially including erroneous bases were removed and clustered based on amplicon sequence variants (ASVs) at single-nucleotide resolution using the DADA2 plug-in (Callahan et al., 2016). The ASVs were taxonomically classified with an amplicon region (the V3-V4 region of 16S rRNA)-specific naïve Bayes classifier derived from 99% OTUs in SILVA rRNA database ver.138.1 using RESCRIPt (Quast et al., 2013; Robeson 2nd et al., 2021). The data were subsequently rarefied to the minimum number of reads (17,024 reads) to normalize the read counts across samples.

The alpha-diversity of the bacterial community among the samples was determined by Pielou’s eveness and Shannon indices, and differences in the diversity between groups except for the seawater sample were statistically analyzed by the Kruskal–Wallis test (q < 0.01) using QIIME2. The beta-diversity of the bacterial community among the groups was determined by the Bray-Curtis index, and the significant differences between groups were analyzed by the PERMANOVA test (q < 0.01) using QIIME2. All the statistical analyses were followed by Benjamini–Hochberg false discovery rate correction. Both QIIME2 and R Studio (v. 2023.09.1.494) with the ggplot2 package were used for data visualization (Bolyen et al., 2019; R Core Team, 2023; Posit team, 2023).

## Results

### Embryo survival rate in the extracapsular environment

Developing catshark embryos were removed from the capsule and reared in a separate container filled with natural seawater with or without antibiotics (Figure 2). Embryos just prior to the pre-hatching period (st. 30-31E) survived for at least 20 days in seawater outside the capsule, regardless of the presence of antibiotics (Figure 2A). Earlier embryos (st. 27-28) survived for 20 days in seawater supplemented with antibiotics, whereas the survival rate dropped to <50% within 4 days in the absence of antibiotics, and all embryos died within 18 days under this condition (Figure 2B).

**Figure 2:**
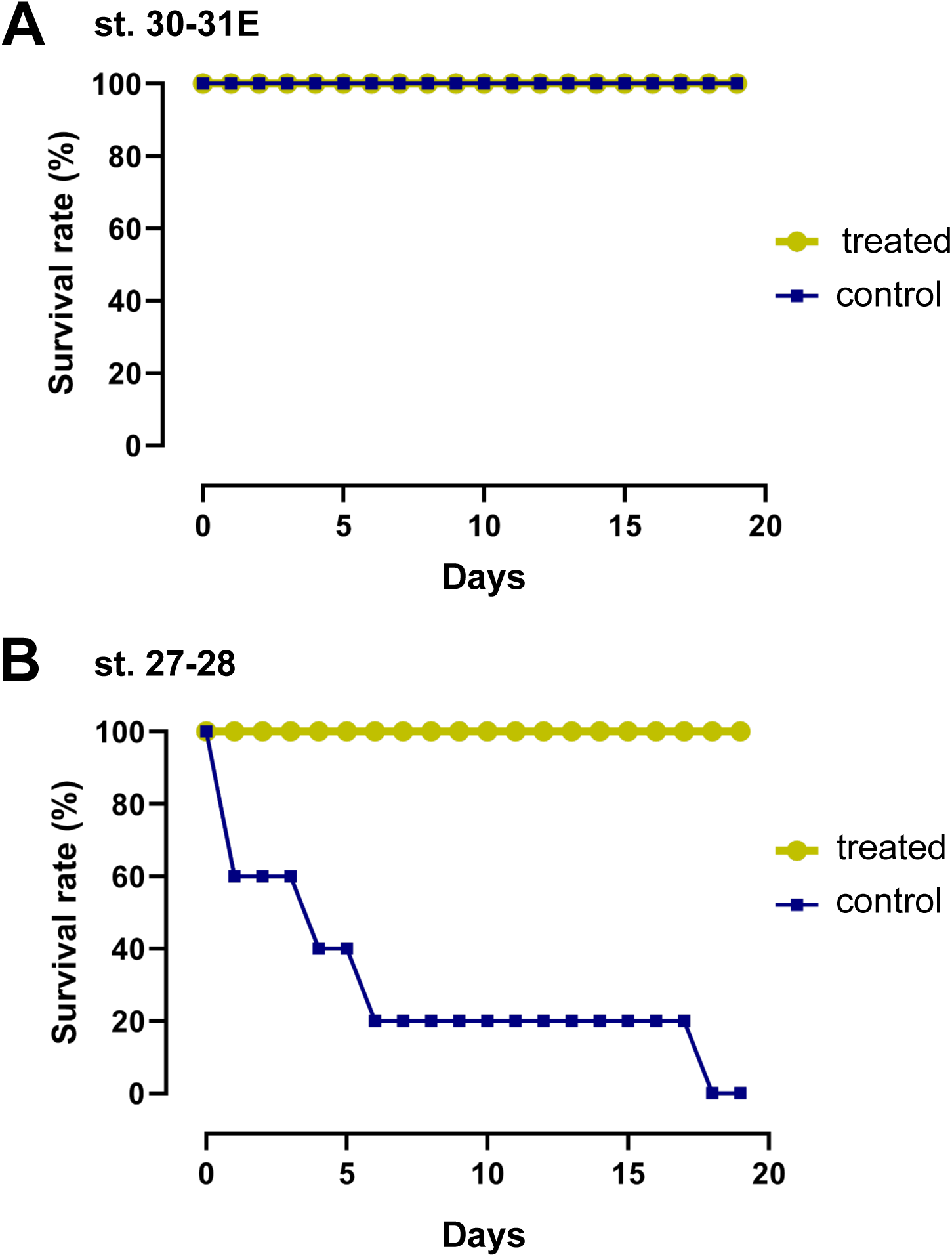
Embryo survival rate of catshark outside the capsule at two different developmental stages. The embryos were reared in natural seawater with (treated) or without (control) antibiotics. **A** Embryos at later stage (st. 30-31E). **B** Embryos at earlier stage (st. 27-28).

### Abundance of microorganisms within the egg capsule

The bacterial density in the internal environment of freshly laid eggs, dead eggs, eggs carrying embryos at two different stages, and seawater was measured by counting the number of DAPI-positive cells (Figure 3A). The density in the jelly of freshly laid eggs was >40 times lower (3.75 × 10^4^ ± 0.86 × 10^4^ cells/mL) than that of seawater in the container rearing the eggs (1.62 × 10^6^ cells/mL). Similarly, low values were obtained for the eggs carrying embryos, although these values were slightly greater than that of freshly laid eggs: st. 27-28, 6.61 × 10^4^ ± 0.98 × 10^4^ cells/mL; st. 30-31E, 6.38 × 10^4^ ± 0.73 × 10^4^ cells/mL. There were no significant differences between the three groups.

**Figure 3:**
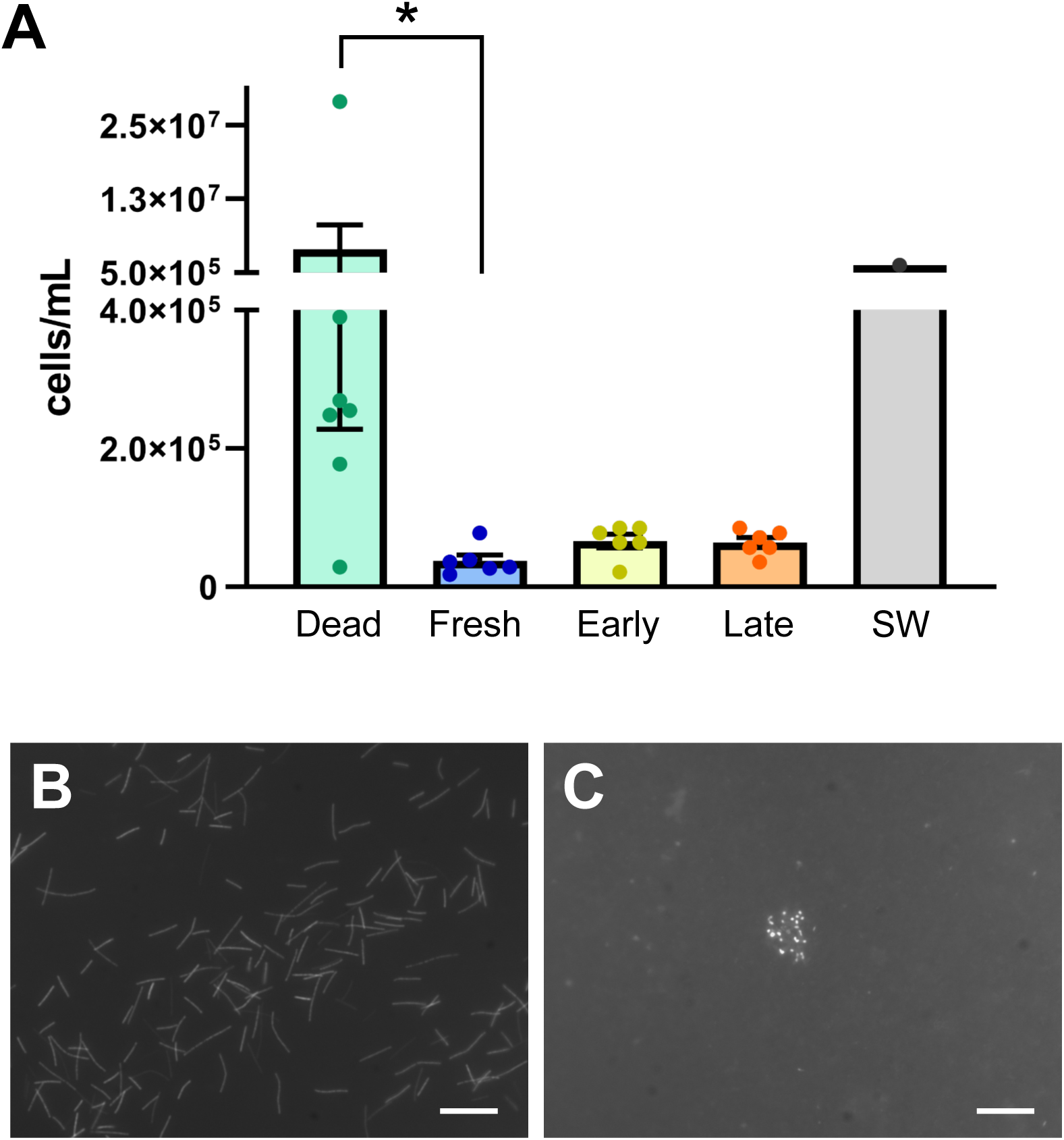
Microbial abundance inside the egg capsule of catshark. **A** Estimated numbers of bacteria per unit volume (cells/mL) in the contents inside the capsule. The egg conditions were as follows: Dead (dead eggs, *N* = 7), Fresh (freshly laid eggs, *N* = 6), Early (eggs carrying embryos at stage 27-28, *N* = 6), and Late (eggs carrying embryos at stage 30-31E, *N* = 6). The values are presented as the means ± SEM. The microbial density of the rearing seawater (SW) was also examined for comparison (*N* = 1). Asterisk indicates the statistical significance (*p* < 0.05). **B** Conspicuous bacterial outgrowth in selected dead eggs. **C** A representative bacterial aggregate found in freshly laid eggs. Scale bars, 10 µm.

In contrast, the highest average microbial density was observed in the intracapsular contents of dead eggs (4.25 × 10^6^ ± 4.02 × 10^6^ cells/mL, similar to that of seawater) (Figure 3A). The bacterial density in dead eggs varied markedly; one dead egg had an extremely low bacterial density, similar to that of freshly laid eggs (2.84 × 10^4^ cells/mL), whereas another dead egg had an exceptionally high density (2.83 × 10^7^ cells/mL), 17 times greater than that of seawater. In the latter case, a large number of rod-shaped bacteria were observed in every observation field (Figure 3B). Interestingly, when the entire filtered membrane was thoroughly observed, we found bacterial aggregates in all samples of freshly laid eggs and eggs carrying embryos, but not in dead eggs or seawater (Figure 3C). The bacteria within the aggregates were not counted.

### Bacterial community within the eggs and OG

All samples of DNA extracted from dead eggs, the OG, and seawater were successfully amplified and sequenced. However, 4 of 10 samples of freshly laid eggs were excluded from the examination due to insufficient PCR amplification for subsequent analyses. The ASV data were rarefied based on the minimum number of reads (17,024) and taxonomically classified into 3,612 features (Supplementary Fig. 1). The assigned features were collapsed to taxonomic level 6, corresponding to “genus” level, and the relative abundance of the top nine taxa was visualized (Figure 4).

**Figure 4:**
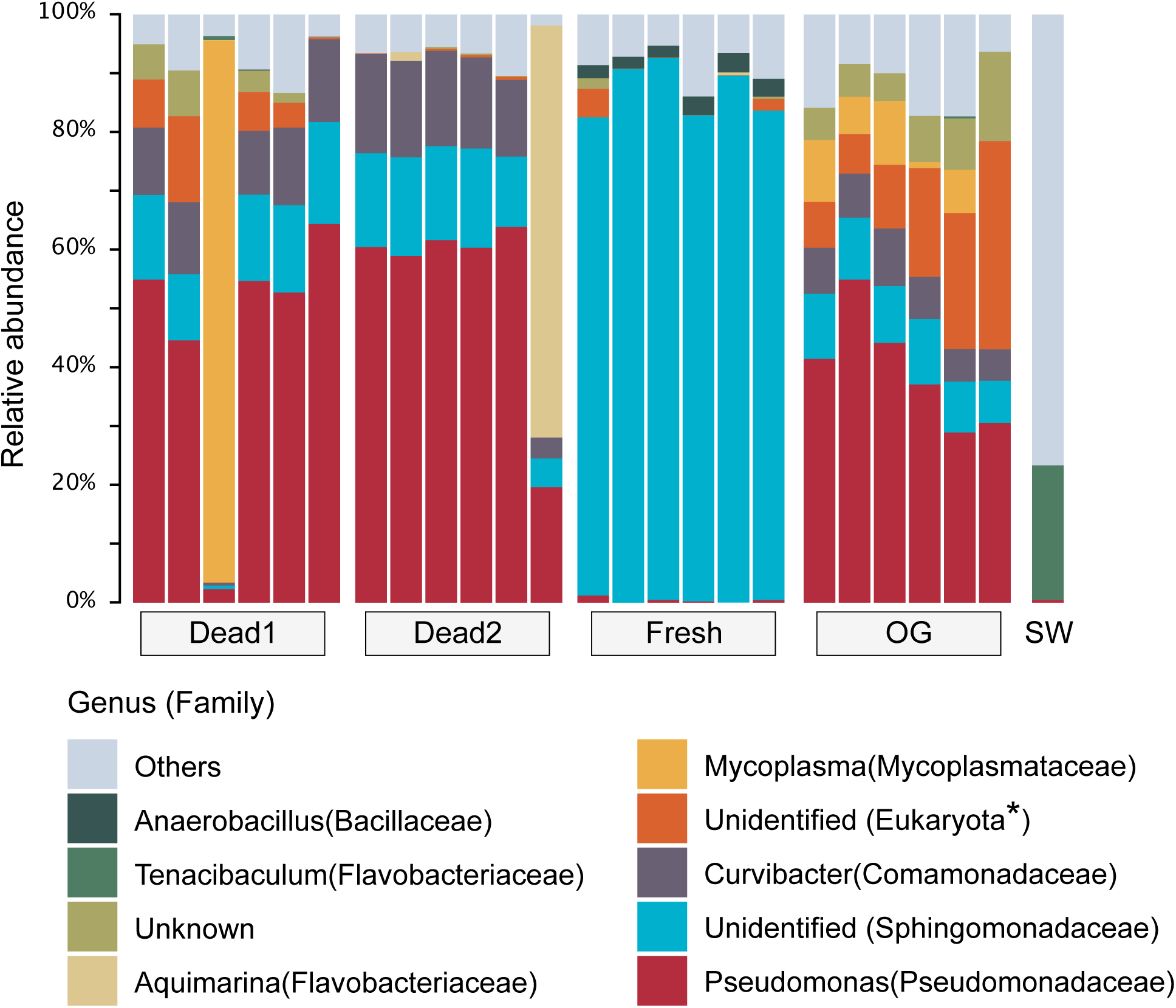
Taxonomic compositions at the genus level. Relative abundance of top 10 ASVs are shown in bar plots. Dead1, dead eggs 1 month after oviposition; Dead2, dead eggs 2.5 months after oviposition; Fresh, freshly laid eggs; OG, epithelia of oviducal gland; SW, seawater. Note that the information in parentheses indicates the family level, with the exception of Eukaryota, which represents the domain level.

In freshly laid eggs, an unidentified genus of the Sphingomonadaceae family (Pseudomonadota phylum, alpha-Proteobacteria class) dominated the microbial community, comprising more than 81.8% in all six examined samples, with the highest sample reaching 92.1% (Figure 4). Although the unidentified Sphingomonadaceae genus was also present in all the other samples, except for seawater, the occupancy was less than 17.4%. Meanwhile, 10 of 12 samples of dead eggs were dominated by *Pseudomonas* (57.6% on average, Proteobacteria phylum, gamma-Proteobacteria class, Pseudomonadaceae family), followed by *Curvibacter* (13.8% on average, Pseudomonadota phylum, gamma-Proteobacteria class, Comamonadaceae family). Furthermore, the bacterial communities in two individual samples of dead eggs were unique and distinct from those in other samples. The community in one dead eggs 1 month after oviposition exhibited a high occupancy of *Mycoplasma* (92.1%), whereas apredominance of *Aquimarina* (70.4%) was found in another dead egg 2.5 months after oviposition.

Similar to the microbiota of dead eggs 1 month after oviposition, the microbiota of the OG was dominated by *Pseudomonas*, *Curvibacter*, and unidentified genera of the Sphingomonadaceae family. Eukaryote ASVs were identified in all OG samples, ranging from 6.5% to 35.4%, with an average of 17.1%. Contamination by eukaryotic nucleotides (8.5% on average) was also observed in four of six dead eggs 1 month after oviposition, but this contamination was <0.6% in dead eggs 2.5 months after oviposition. These eukaryotic ASVs in catshark-derived samples were top-hit-matched to the *S. canicula* genomic sequence based on a BLASTn search, confirming host origin. In contrast to the communities in the catshark-derived samples, the microbial community in the seawater sample was highly diverse and primarily dominated by *Tenacibaculum* (22.8%, Bacteroidota phylum, Bacteroidia class, Flavobacteriaceae family), followed by various taxa at <10% abundance.

### Bacterial diversity

When analyzing alpha-diversity using both the Shannon and Pielou indices, freshly laid eggs showed the least diversity, whereas seawater exhibited the greatest diversity (Figure 5A, B). According to both the Shannon and Pielou indices, the alpha-diversity of freshly laid eggs differed significantly from that of the OG and dead eggs, consistent with the results of the community analyses (Figures 4 and 5A, B). The OG exhibited the greatest diversity among catshark-derived samples and differed significantly from that of dead eggs 2.5 months after oviposition. However, there was no differences in the diversity between the OG and dead eggs 1 month after oviposition. No significant differences were detected between the two groups of dead eggs (i.e., eggs of different postmortem durations).

**Figure 5:**
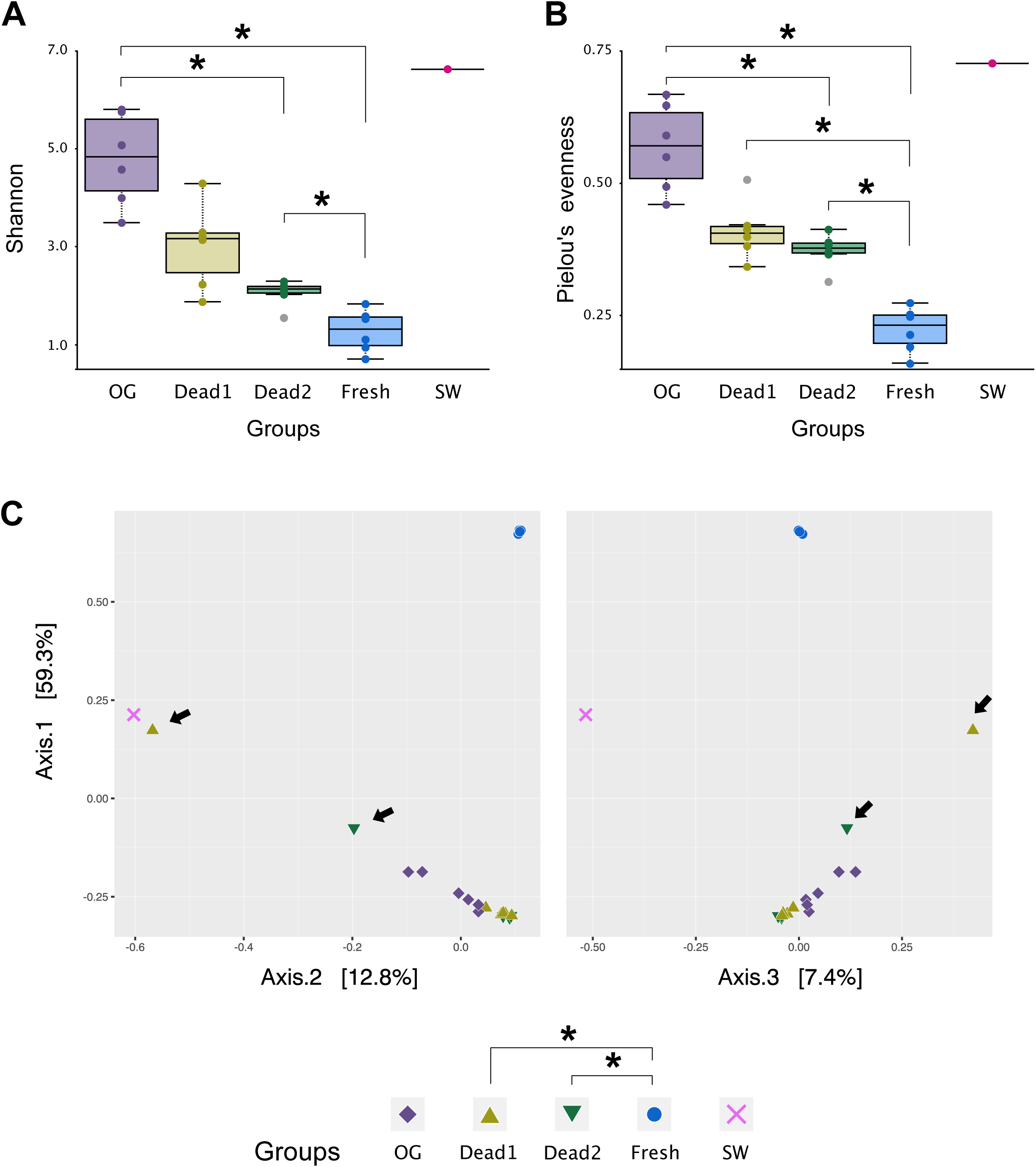
Results of alpha- and beta-diversity. Asterisks indicate the statistical significance (q < 0.05). The abbreviation is the same as those in Figure 4. **A** Shannon and **B** Pielou’s evenness were used for calculation of alpha-diversity. Outlier samples are indicated by gray circles. **C** The Bray-Curtis dissimilarity index was used for beta-diversity analysis. The PCoA plot shows the relatedness of microbial community among the examined groups. Arrows indicate two dead egg samples dominated by *Mycoplasma* and *Aquimarina*, respectively.

Taxonomic beta-diversity was calculated using Bray-Curtis dissimilarity analysis, and the results are represented on principal coordinate axes (Figure 5C). All samples of freshly laid eggs clustered closely and showed low similarity with the other groups. Consistent with this result, statistical analysis revealed significant differences between freshly laid eggs and dead eggs. However, no significant difference in taxonomic diversity was observed between the OG and other samples. The two samples of dead eggs that exhibited unique bacterial compositions in the previous community analysis showed variability in spatial position (Figure 4, arrows in Figure 5C).

## Discussion

### Embryonic immune functions in catshark

The inability of early embryos of oviparous elasmobranchs to survive outside the egg capsule has long been known (Ballard et al., 1993), and the addition of antibiotics to the rearing seawater is a well-established empirical viability-extending treatment (Onimaru et al., 2018; Kuroda et al., 2021). However, the point at which embryonic defense against pathogens is acquired during development has not been experimentally verified. In cloudy catshark, differences in the ability of early embryos to survive outside the egg capsule were observed between two developmental stages (st.27-28 and st.30-31E). This discrepancy is likely attributed to differences in embryonic resistance to potential pathogens, as all examined embryos survived with antibiotic treatment. Previous studies examining *S. torazame* demonstrated that the pre-hatching period (st.31) is a critical time during which embryos develop respiratory, osmoregulatory, and nutritional absorption functions (Tomita et al., 2014; Takagi et al., 2017; Honda et al., 2020). It is thus reasonable to consider that the period around developmental stage 31 is also a key time for the acquisition of immune functions.

The transfer of passive immunity from mothers to embryos, in which maternal antibodies accumulated in egg yolk contribute to embryonic defense, has been documented in a variety of vertebrates (Mor and Avtalion, 1990; Olsen and Press, 1997; Hamal et al., 2006). For example, the presence of maternally derived immunoglobulins (7S IgM and IgNAR) in the egg yolk of viviparous nurse sharks (*Ginglymostoma cirratum*) suggests these immunoglobulins contribute to embryonic immunity (Haines et al., 2005). However, the inability of young catshark embryos at stages 27-28 to survive outside the capsule without antibiotics suggests the maternal contribution to embryonic defense in catshark is less significant, although whether maternal antibodies are present in the eggs is currently unknown. Our data support the conclusion that early-stage embryos of oviparous elasmobranchs lack resistance to pathogenic bacteria.

### Low microbial density within the egg capsule and its maintenance

Mika et al. (2021) characterized microbial communities in embryonic tissues, egg capsules, and jelly in the oviparous little skate and demonsted the vertical transfer of the microbiome from mother to embryo. However, the absolute bacterial density in oviparous elasmobranchs has not been reported. Considering the results of the survival test, it is evident that a clean intracapsular environment is necessary for survival of immunologically immature early embryos. Indeed, the jelly inside freshly laid eggs of catshark exhibited low bacterial density, which was maintained until pre-hatching, and the bacterial community was unique compared with that of other samples, with low species richness. The intracapsular fluid of little skate at stage 0 (equivalent to the freshly laid egg in this study) and stage 16 showed very low species richness compared with later stages (Mika et al., 2021), in good accordance with our findings. Thus, a clean intracapsular environment for eggs at oviposition might be common among oviparous elasmobranchs.

A tertiary egg coat that functions as a primary barrier for preventing the invasion of bacteria from the external environment is common among oviparous vertebrates. In chicken and Chinese soft-shell turtle, the pore sizes of the membrane lining the eggshell is reportedly 25.2 and 20 nm, respectively (Kutchai and Steen, 1971; Yoshizaki et al., 2004). The chorion in teleost fishes also exhibits a submicron pore size, which is smaller than typical micron-scale bacteria (Olivar, 1987). Likewise, the calculated pore size of the egg capsule wall in *S. canicula*, a close relative of *S. torazame*, is several nanometers (Hornsey, 1978). Considering these data together with our finding that the microbial compositions of both freshly laid eggs and dead eggs were distinct from that of seawater confirms that the catshark egg capsule wall functions as a structural barrier against the entry of microorganisms and thus likely contributes to the maintenance of a closed environment before pre-hatching.

#### Mechanism leading to generation of the unique intracapsular environment

Determining the mechanism by which the clean intracapsular environment is generated remains challenging. The aseptic condition within freshly laid eggs is a well-known characteristic of avian species. Various physicochemical properties (e.g., temperature, pH, and high viscosity) and an abundance of antimicrobial proteins (such as ovo-transferrin and lysozyme) in the egg white of chicken inhibit bacterial growth (Alabdeh et al., 2011; Giansanti et al., 2012; Baron et al., 2016). The presence of lysozyme in egg white has also been reported in reptiles (Gayen et al., 1977). In contrast, the egg jelly of oviparous elasmobranchs is a mucin hydrogel containing a substantial amount of water with a very low protein level (<0.001%; Lenain and Henderson, 2020). Lenain and Henderson (2020) further demonstrated that the jelly does not affect the growth of bacteria isolated from the outer surface of the egg capsule, negating the contribution of the egg jelly to embryonic defense. However, it is not necessary that the jelly, which has no contact with bacteria outside the capsule, exhibit strong antibacterial activity. Indeed, the egg jelly from *Scyliorhinus stellaris* shows a slight inhibitory effect against gram-positive bacteria (Martinengo et al., 2024). Thus, the jelly appears to play a bacteriostatic role in early development, given that the initial lowered bacterial abundance at the oviposition was maintained until pre-hatching.

Within freshly laid eggs and eggs carrying developing embryos at two different stages, we observed bacterial aggregates, albeit infrequently, with the numbers remaining relatively stable throughout the examined developmental periods. The formation of bacterial aggregates has generally been reported as a response to infection, and a variety of factors mediate the induction of aggregation (Secor et al., 2018; Cai, 2020). In humans, for example, interactions between IgA secreted in saliva and bacterial surface components induce aggregation to promote clearance by blocking bacterial adhesion to oral tissue surfaces (Yamaguchi, 2004). Similarly, mucus secreted by the colon induces the aggregation of intestinal bacteria, preventing inflammation (Bergström et al., 2016). In addition, environments rich in polymers such as mucins and glycosaminoglycans also drive bacterial aggregation via entropic forces (Secor et al., 2018; 2022). Therefore, the formation of aggregates found inside the capsule might be induced by the mucin hydrogel and its degradation products, thereby suppressing bacterial activity and/or proliferation before pre-hatching. Further research is warranted to explore the potential bacteriostatic action of the jelly.

A potential contributor to reduction in intracapsular bacterial biomass before oviposition is the oviducal gland, the site of both fertilization and encapsulation in elasmobranchs (Wourms, 1977; Hamlett et al., 1998). As the bacterial flora inside freshly laid eggs differed significantly from that of oviducal gland epithelia, it is plausible that bactericidal substances are secreted from the oviducal gland to sterilize the luminal epithelium, thereby minimizing pathogen contamination of the capsule during the process of encapsulation.

In dead eggs, nutrient-rich yolk contents diffuse within the capsule, potentially serving as a nutrient source that triggers explosive bacterial growth. We identified potential pathogenic bacteria, specifically *Mycoplasma* and *Aquimarina*, in two dead eggs. Given the well-known pathogenicity of species belonging to these genera in eukaryotes (Razin et al., 1998; Hudson and Egan, 2022), these bacteria could also be harmful to catshark embryos. Notably, *Mycoplasma*, which dominated the oviducal gland to a certain extent, was not detected in other egg samples. This finding suggests the gland has a mechanism for selectively eliminating pathogenic bacteria from the eggs.

Production of bactericidal or bacteriostatic substances in adult reproductive organs is well documented among vertebrates (King et al., 2003; Yoshimura and Barua, 2017; Iida et al., 2021). Recent transcriptomic and proteomic analyses of the uterine fluid of viviparous red stingray (*Hemitrygon akajei*) revealed the presence of antimicrobial transferrins and dermicidin throughout the examined gestational periods (Kina et al., 2021). It should be noted that the adult female animals dissected in this study were not in the state of forming egg capsules due to difficulties in identifying the timing of fertilization and subsequent egg encapsulation. Thus, the use of plasma progesterone concentration as an indicator of ovulation (Inoue et al., 2020; Shimoyama et al., 2023) might allow us to obtain females just at the time of egg encapsulation. Examining the microbial community of the oviducal gland in these animals would provide insights into the mechanism by which the number of bacteria inside the capsule is reduced before oviposition.

### Relationship between the microbiota and embryonic growth

As in other vertebrates, interactions between the host and microbes have been predicted in elasmobranchs (Perry et al., 2021). In catshark, the freshly laid eggs, almost completely dominated by a single unidentified genus of Sphingomonadaceae, exhibited a highly unique bacterial community distinct from that of other examined samples. Bacterium belonging to the Sphingomonadaceae was also found in the oviducal gland to a certain extent, but it was undetectable in seawater, suggesting the occurrence of transgenerational microbial transfer from mother to fetus, as reported in little skate (Mika et al., 2021). The notably high occupancy of this bacterium in the freshly laid eggs raises the possibility of a symbiotic relationship with the host, although the benefits of this relationship remain to be determined. Several genera belonging to the family Sphingomonadaceae contain carotenoids that are known to act as antimicrobial agents (Kosako et al., 2000; Vargas-Sinisterra and Ramírez-Castrillón 2021). Thus, a carotenoid-mediated antimicrobial mechanism may be active in the freshly laid eggs of catshark.

## Conclusions

This study provides the first direct evidence that early embryos of oviparous cloudy catshark are incubated in a clean intracapsular environment that minimizes the number of microorganisms until pre-hatching. The intracapsular environment of freshly laid eggs in this study exhibited extremely low microbial diversity and was largely dominated by an unidentified genus of Sphingomonadaceae. Our findings enhance understanding of the mechanism of embryonic protection against pathogens and may provide a novel approach for the exploration of microbial associations with elasmobranch embryos.

## Acknowledgements

We are grateful to the staff of Oarai Aquarium for their assistance in obtaining adult catshark. Our sincere gratitude is extended to Dr. Koji Hamasaki, Dr. Takuhei Shiozaki, and their laboratory members for providing laboratory space and equipment.

Computations were partially performed on the NIG supercomputer at ROIS National Institute of Genetics.

## Authors’ contributions

WT conceived the project, designed the experiments and wrote the original draft. AM, YS-T, KS, KT, and WT contributed to sample collection and the preparation. AM, YS-T, and WT performed the experiments. All authors participated in data analysis and discussions during the manuscript preparation. All authors approved the final version of the manuscript.

## Ethics declarations

### Ethics approval and consent to participate

All animal experiments were approved by the Animal Ethics Committee of Atmosphere and Ocean Research Institute of the University of Tokyo (P19-2). The present study was carried out in compliance with the ARRIVE guidelines.

### Consent for publication

Not applicable.

