## Supplementary figures and images for "Low microbial abundance and community diversity within the egg capsule of the oviparous cloudy catshark (*Scyliorhinus torazame*) during oviposition"

### Supplementary Figure 1

#### Supplementary Figure 1

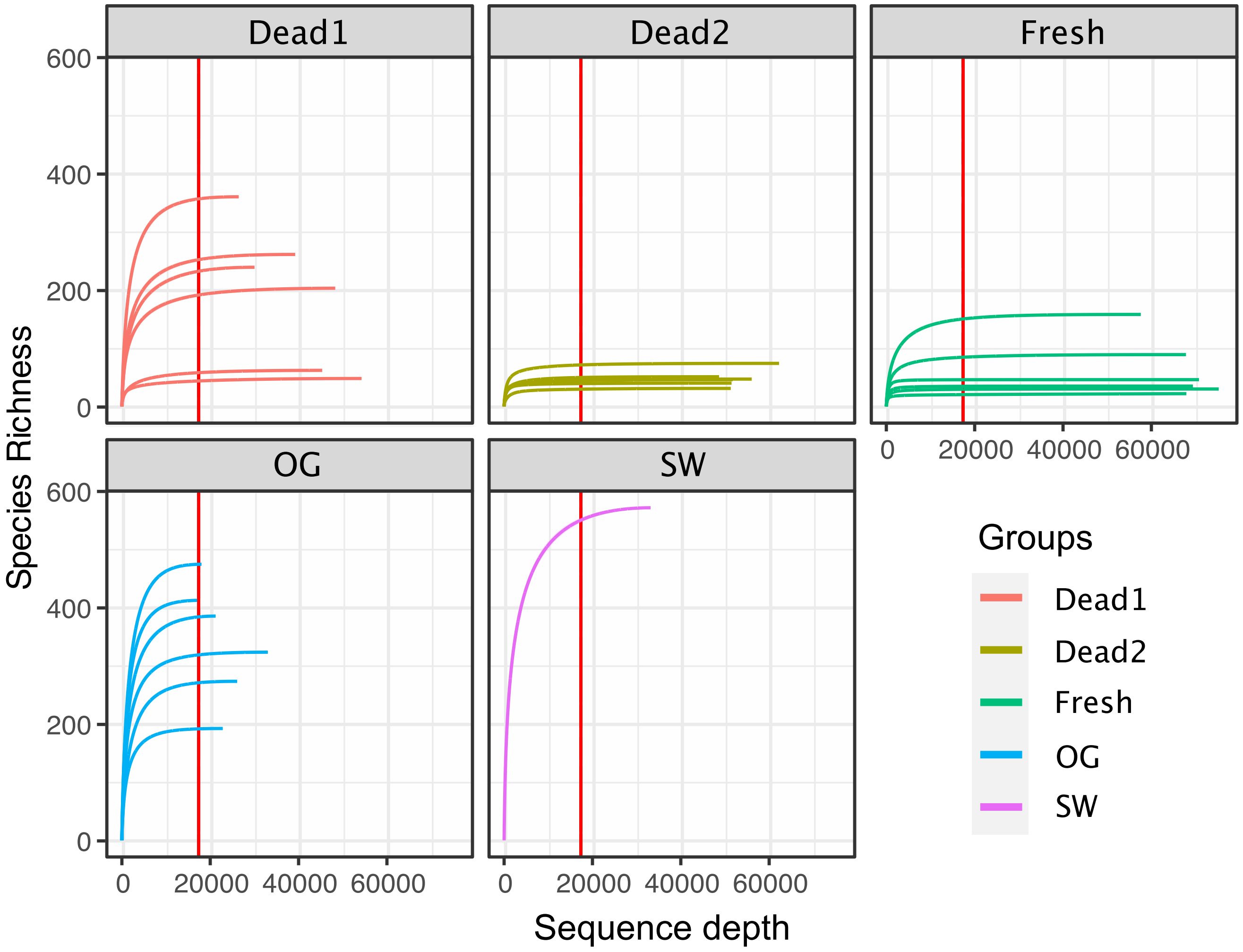
