## Supplementary Figure Legend for "Low microbial abundance and community diversity within the egg capsule of the oviparous cloudy catshark (*Scyliorhinus torazame*) during oviposition"

**Supplementary Figure 1**: Alpha rarefaction curve of the samples. Red lines indicate the read count of the sample with the minimum number of reads (17,024 reads). The abbreviation is the same as those in Figure 4.
